# Abnormal craniofacial and spinal bone development with *col2a1*a depletion in a zebrafish model of CHARGE syndrome

**DOI:** 10.1101/2020.07.10.197533

**Authors:** Maximilian Breuer, Maximilian Rummler, Charlotte Zaouter, Bettina M. Willie, Shunmoogum A. Patten

## Abstract

CHARGE syndrome patients commonly display craniofacial abnormalities. Furthermore, most patients show features of idiopathic scoliosis, reduced bone mineral density and in a few cases osteopenia. While several clinical cases and studies have documented the skeletal deformities in CHARGE syndrome bearing *CHD7* mutations, the underlying mechanisms of the disorder remain elusive. Here, we detect and quantitatively analyze skeletal abnormalities in larval and adult *chd7*^-/-^ zebrafish.

We show that young *chd7*^-/-^ larvae present with abnormal craniofacial development, especially related to cartilage. We also observe scoliosis-like spinal deformations at 9 dpf. Gene expression analysis confirmed the reduction of osteoblast markers and Pparγ targets. MicroCT analyses identified abnormal craniofacial structures, Weberian apparatus and vertebral body morphology in *chd7*^-/-^ mutants, with highly mineralized inclusions, along with significant variances in bone mineral density and bone volume. Notably, we detect a specific depletion of Col2a1a in the cartilage of craniofacial regions and vertebrae, in line with a significantly reduced number of chondrocytes.

Our study is the first to elucidate the mechanisms underlying morphological changes in craniofacial structure and vertebrae of adult *chd7*^-/-^ zebrafish. The *chd7*^-/-^ mutant zebrafish will be beneficial in future investigations of the underlying pathways of both craniofacial and spinal deformities commonly seen in CHARGE syndrome.

## Introduction

Impaired bone development, craniofacial dysmorphism and spinal abnormalities are a major health concern in various genetic disorders. Among these, idiopathic scoliosis is the most common skeletal abnormality observed in children (1). Idiopathic scoliosis is a curvature of the spine with unidentified genetic cause. However, many risk genes have been connected to the underlying mechanisms (2). One of these related genetic disorders is the congenital multisystemic CHARGE syndrome (CS) named after the major characteristics of **C**oloboma, **H**eart defects, **A**tresea chonae, **R**etarded growth, **G**enital and **E**ar abnormalities (3–5). Though, considering the wide variety of observed phenotypes, the list of characteristics has been continuously augmented to include craniofacial abnormalities and spinal deformations (6–8). Along these lines, CS is closely associated with skeletal deformities such as craniofacial dysmorphisms, idiopathic scoliosis, kyphosis and hemivertebrae (9). In fact, idiopathic scoliosis is observed in a majority of CS cases, with studies showing over 60% of patients having diagnosed scoliosis at an average age of just over 6 years (10–12). In some cases, the areal bone mineral density (aBMD) is reduced (13).

CS is most commonly caused by a mutation in the chromodomain ATP-dependant helicase 7 (CHD7). Mutations are distributed evenly throughout the gene and the vast majority of cases are sporadic, with only very few cases of familial CS (7, 14–16). Analysis of pathways related to CHD7 show involvement in neural crest differentiation/proliferation/migration and stem cell quiescence (17–19). Furthermore, studies focusing on CS have identified regulatory mechanisms of CHD7 in the immune response and more strikingly in brain development (20, 21). Most recently, CHD7 has been connected to abnormal GABAergic development resulting in an autistic-like behavior (Jamadagni et al., unpublished data). While the function of CHD7 has been extensively studied, little attention has been given to the underlying mechanisms in skeletal development (22). Some studies have directly linked CHD7 to osteogenesis (23, 24).

Investigation in cell cultures have shown a dependency on CHD7 for successful differentiation, more specifically to a differentiation complex with SETD1B, NLK and SMAD1 resulting in depleted osteogenesis upon depletion of CHD7 in favour of adipogenesis by regulating PPARγ target genes (23, 24). Various models to investigate this role in vivo have been proposed. A mouse model for Chd7 deficiency, termed “looper” presents with ear ossicle malformations, but no spinal deformities (25). However, mouse models for Chd7 deficiency have limitations, as null mutants are embryonically lethal by 10 days (26). Yet, a recent zebrafish model has proven valuable in the modelling of *chd7* dependent CS revealing a reduction in vertebrae mineralization of young larvae (27).

Zebrafish have become increasingly relevant in the study of fundamental bone development and bone related disorders (28–30), including scoliosis, osteoporosis, age related osteoarthritis (31, 32). Notably, the spinal structure in zebrafish has been characterized in detail and comprises three major regions: the Weberian apparatus, a specialized structure consisting of the first four vertebrae connecting the auditory system and the swim bladder to amplify sound vibrations, the precaudal vertebrae which are connected to neural arch and spines and caudal vertebrae of the tail region with neural and hemal arch (33). Simplicity of analysis in zebrafish to investigate spinal structures has been used to understand the effects of mechanical loading on bone mineralization (30, 34). Zebrafish have specific advantages in the analysis of skeletal development such as the closely related structure to humans. They also have rapid bone development with first mineralization of vertebrate occurring after only 5 days post fertilization (dpf). Additionally, the *ex utero* development and transparency of young larvae simplifies the use of *in vivo* staining and transgenic lines to allow for the analysis of early calcification in zebrafish bony structures (35–38). Furthermore, the simplicity of high-throughput drug screening in this teleost model makes the model highly useful for investigating new drug targets and effect on skeletal development (39, 40). These advantages along with their high remodelling efficiency of skeletal structures including their possibility to regenerate fins, as well as the effectiveness to reproduce spinal deformity phenotypes have raised interest in the zebrafish model (41, 42).Studies in zebrafish have shown a close link to collagen related genes (43, 44). These highly conserved pathways between zebrafish and humans allow us to investigate the underlying pathomechanisms linking CHD7 to idiopathic scoliosis and other skeletal abnormalities.

In the present study we describe for the first time a detailed analysis of craniofacial and skeletal anomalies in a zebrafish *chd7*^-/-^ mutant model for CS. Early larvae screens reveal a striking dysmorphism of craniofacial structures, as well as delay of mineralization of vertebrae bodies linked to deficient osteoblast differentiation. Gene expression and IHC analysis reveals striking reduction in osteoblast, chondrocyte, and collagen matrix markers, particularly displaying abolished levels of Col2a1. Finally, extensive microCT analysis in adults show abnormal mineralization and morphology in all major structures of the zebrafish skeleton.

## Results

### Zebrafish chd7^-/-^ larvae show craniofacial and spinal deformities

We recently generate a *chd7* knockout zebrafish line using CRISPR/Cas9 with a single nucleotide insertion causing a frame-shifting mutation and a premature stop codon 8 amino acids after the mutation site (Jamadagni et al., unpublished data). Consistent with our previous findings using a transient *chd7* morpholino knockdown model (27), this stable *chd7*^-/-^ zebrafish mutant using CRISPR/Cas9 that replicates hallmarks of CS including craniofacial and skeletal defects (Fig. 1). To further investigate the bone deformities in *chd7*^-/-^ zebrafish in detail, we first screened young *chd7*^-/-^ larvae at 6 dpf and 9 dpf for morphological abnormalities at early developmental stages. At 6 dpf we observed craniofacial changes, such as reduced length of the palatoquadrate and increased angle of the ceratohyal (Fig 1A). Calcein staining also reveals reduced mineralization of the facial structures, noticeably towards the quadrate and opercle (Fig 1B). At this stage *chd7*^-/-^ larvae were minimally, yet significantly, smaller than control (Cntrl) larvae, however this is in line with the expected CS phenotype of growth retardation and neural crest abnormalities in patients (Supplemental Fig. 1A).

**Fig 1.**
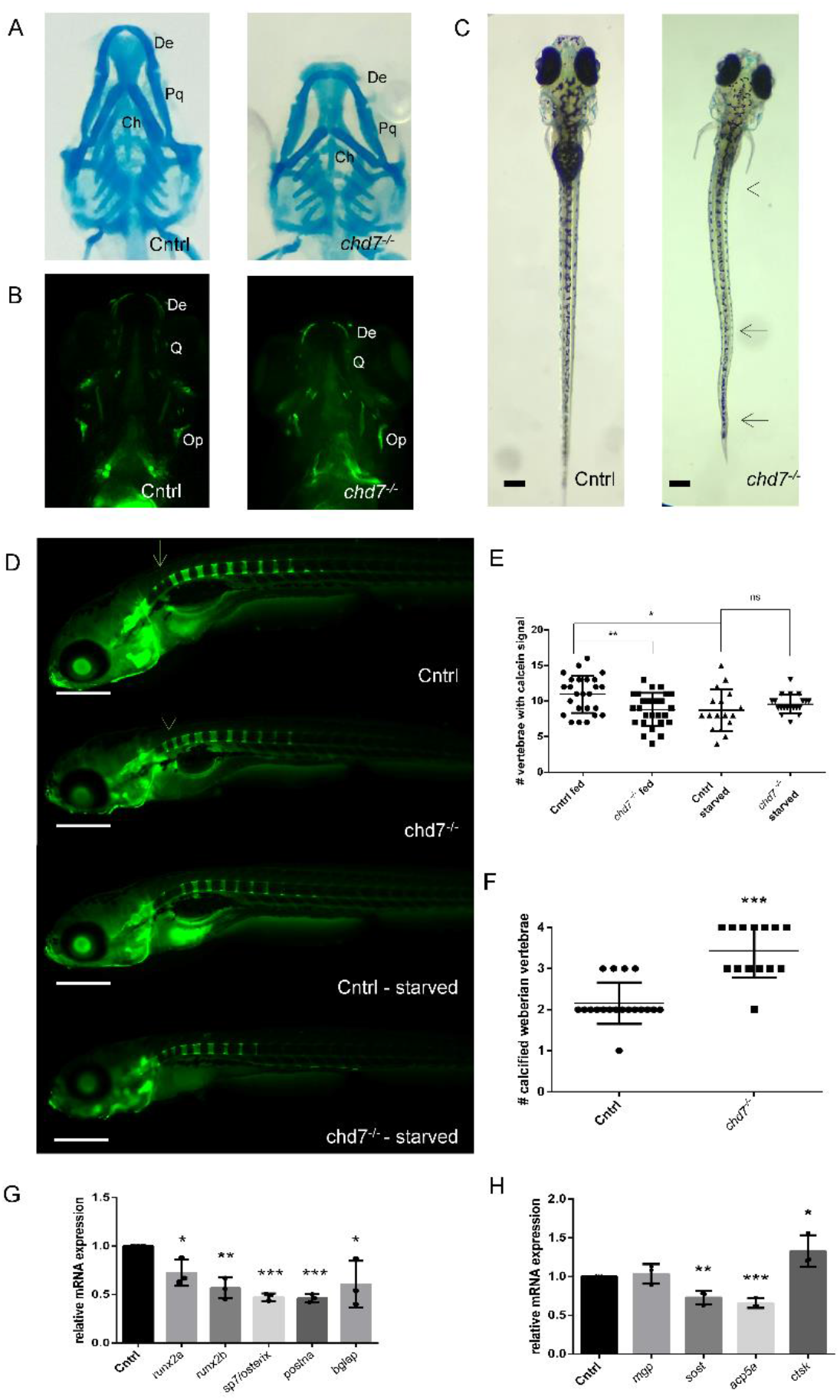
Mineralization in larval *chd7*^-/-^ mutants. A. Alcian blue staining of 6 dpf control and *chd7*^-/-^ zebrafish larvae, ventral view (De: dentary; Pq: palatoquadrate; Ch: ceratohyal) B. Calcein staining of 6 dpf control and *chd7*^-/-^ zebrafish larvae, ventral view (De: dentary; Q: quadrate; Op: opercle) C. Dorsal view of control and *chd7*^-/-^ larvae at 9 dpf showing severe spinal deformities in 43 out of 110 larvae screened. Deformities were present at both precaudal and caudal region (arrows). D. Lateral view of calcein staining at 9 dpf showing mineralization of vertebrae with access to food starting at 5dpf and calcein staining at 9 dpf during starvation. Arrow indicates mineralization at the first 4 (Weberian) vertebrae E. Number of vertebrae in wildtype and *chd7*^-/-^ (fed and starved) larvae showing calcification signal as tested by calcein and F. Number of Weberian vertebrae (vertebrae 1-4) showing complete calcification (see arrows in b) G-H. RT-qPCR of osteogenesis related genes screened at 9dpf, showing relative fold change. (Significance: *p<0.05; **p<0.01; ***p<0.001)

39% of *chd7*^-/-^ larvae exhibited highly varying, scoliosis-like phenotypes at both precaudal and caudal regions of the spine (Fig. 1C). Additionally, some of these also presented with a kyphosis-like phenotype (Supplemental Fig. 1B). We found that *chd7*^-/-^ larvae were still reactive to a touch response, even with the severe morphological phenotype.

### chd7^-/-^ larvae have a spinal mineralization deficit that is nutrient dependent

Given the spinal deformities in 9 dpf *chd7*^-/-^ larvae, we next sought to investigate mineralization during larval development. Calcein staining for fluorochrome labeling of bones revealed no delay in the onset of mineralization of the first precaudal vertebrae at 6 dpf. Upon further development, at 9 dpf, *chd7*^-/-^ larvae showed a significant reduction in the number of calcified vertebrae (Fig. 1D, E). Notably, *chd7*^-/-^ larvae were less efficient in complete calcification of vertebrae towards the posterior region of the spine (Fig. 1D).

Since calcification of the bone matrix is dependent on mineral uptake from food, we tested how the onset and development of vertebral bone structure calcification in 9 dpf *chd7*^-/-^ zebrafish was dependent on nutrition accessibility. Notably, wildtype and *chd7*^-/-^ larvae showed no difference in the number of mineralized vertebrae in a starvation situation between 5 dpf and 9 dpf. Expectedly, we observed significant differences when food was available from 5 dpf until 9 dpf, with *chd7*^-/-^ zebrafish having a reduced number of mineralized vertebrae compared to controls. The *chd7*^-/-^ larvae had the same number of mineralized vertebrae when food was available or absent (Fig. 1 D, E). Starvation in zebrafish larvae between 5 dpf and 9 dpf also resulted in smaller yolks, which is to be expected, considering the high dependency on nutrition supply by the yolk. Surprisingly, we found that Weberian vertebrae, which are the first 4 vertebrae of the spinal column, were more rapidly calcified in *chd7*^-/-^ larvae than in controls (Fig. 1D arrows, Fig 1 F).

We investigated if mineralization recovered in later development. Reduced mineralization remained evident in 4-week-old juvenile zebrafish with spinal deformities. Alizarin red staining revealed decreased mineralization of caudal vertebrae, with striking deficiency toward the vertebral body (Supplemental Fig. 1. D).

### Osteoblast differentiation is impaired in chd7^-/-^ larvae

CHD7 has been shown to regulate osteogenesis by controlling PPARγ to promote skeletal development and regulate expression of promoting genes. To examine if the Pparγ pathway is directly involved in the observed bone mineralization deficiency and abnormal morphology in *chd7*^-/-^ larvae, we tested for Pparγ target gene expression and osteoblast differentiation marker genes at the onset of the bone defect at 9 dpf (Fig. 1G). The key targets for Pparγ, *runx2a* and *runx2b*, which are expressed mostly in craniofacial regions at this stage, were significantly downregulated in *chd7*^-/-^ larvae compared to controls. Additionally, other markers for osteogenesis were also affected, including a significant downregulation of osteoblast markers *sp7/osterix, bglap* and *postna* (Fig. 1G). The *chd7*^-/-^ larvae also had significantly lower levels of osteocyte markers while other genes regulating mineralization such as *acp5a* and *sost*, while *mgp* remained unchanged compared to control larvae (Fig 1H). We also detected a significant upregulation of the osteoclast marker *ctsk* in *chd7*^-/-^ larvae.

### Abnormal development of craniofacial regions and the Weberian apparatus in adult chd7^-/-^ zebrafish

To assess the effects of *chd7* deficiency on bone development, including craniofacial and vertebral growth and maturation, we analyzed bone morphology and mass using microCT in adult zebrafish. Morphologically, adult *chd7*^-/-^ zebrafish were unchanged in body length.

Since mineralization and morphology in the young larvae varied in craniofacial regions, as well as from anterior to posterior regions of the spine, we decided to analyze the four major bony structures of the zebrafish: the skull, Weberian apparatus, precaudal and caudal vertebrae. The skull was notably changed in *chd7*^-/-^ zebrafish in comparison to controls (Fig. 2A-D). In particular, we observed significant alterations in the mandibular angle and length (Fig. 2E, F). The face of *chd7*^-/-^ zebrafish was also notably deformed compared to control zebrafish, with a significantly increased craniofacial angle (Fig. 2G). Further, the angle of the mandibular arch was wider in *chd7*^-/-^ zebrafish. Lastly, 3 out of 5 *chd7*^-/-^ zebrafish presented with a warped skull, with the tip of the dentary deviating from the fish’s midline.

**Fig 2.**
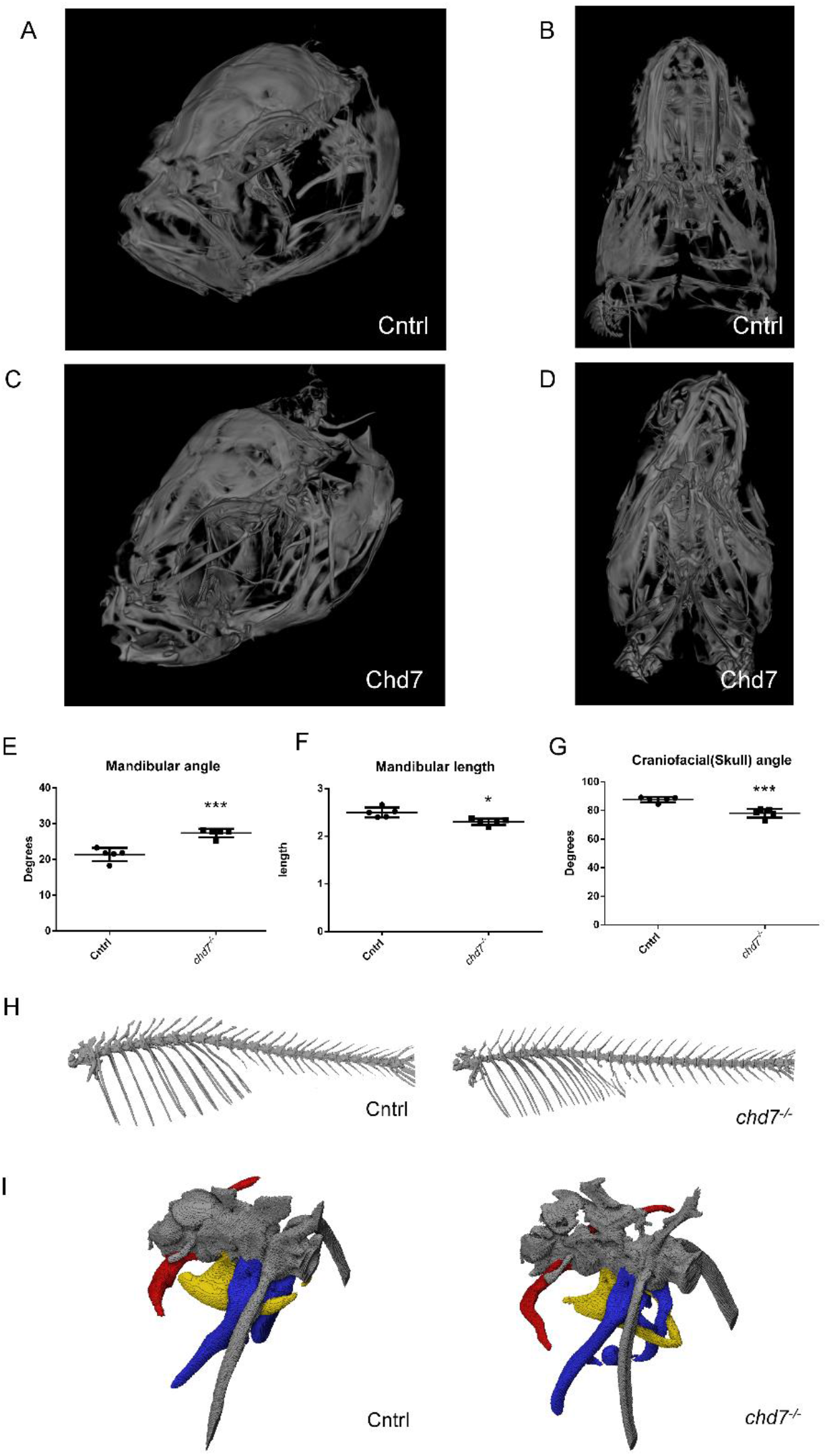
Skull deformities and Weberian apparatus. A, C. Lateral view of Control and *chd7*^-/-^ adult zebrafish skull and B, D. ventral view of Control and *chd7*^-/-^ adult zebrafish of mineralized tissue E. Angle of mandibular arch in adult zebrafish F. Length (in mm) of mandibular bone and G. Angle of the craniofacial (skull) structure H. Representative microCT overview image of the spinal cord from a 1-year old control and a *chd7*^-/-^ adult zebrafish I. MicroCT Image of the Weberian apparatus from a 1-year old control and a *chd7*^-/-^ mutants, indicating structures of intercalarium (red), tripus (yellow) and parapophysis (blue). Significance of student t-test with Welch’s correction is included in the graphs with: *p<0.05; **p<0.01, ***p<0.001

Another key structure, the Weberian apparatus with its supraneurals, intercalarium, tripus and parapophysis, had thinner and smaller morphology in *chd7*^-/-^ adults compared to controls (Fig. 2I). We further performed a detailed analysis of volumetric bone mineral density (vBMD), bone volume (BV), total volume (TV), and bone volume fraction (BV/TV) of 1-year-old *chd7*^-/-^ and control zebrafish. We did not detect any significant differences in vBMD, BV or TV (Supplemental. Fig. 2). However, we found a significantly greater variance in TV of both intercalarium and parapophysis, as well as in BV of the parapophysis in 1-year-old *chd7*^-/-^ zebrafish, as determined by F-test analysis (Supplemental Table 1). The same results were observed in 2-year-old *chd7*^-/-^ compared to control zebrafish (Supplemental Table 2).

### Abnormal mineralization and malformations in the body end plates of precaudal vertebrae in adult zebrafish

Vertebrae structures were analyzed by assessing key angles of the structure (Fig. 3A) and mineralization features. MicroCT analysis of precaudal vertebrae identified abnormal patterns of mineralization and abnormal structures, that were especially pronounced at the arches and transverse processes in 1-year old *chd7*^-/-^ and control zebrafish (Fig. 3B, C). Precaudal vertebrae showed significantly higher overall vBMD, while the distribution of mineralization between the body and arch remained unaffected (Fig. 3D, E). However, structurally, precaudal vertebrae of *chd7*^-/-^ mutants show an increase in bone volume towards the body end plates of vertebrae, which in most cases include major malformations manifesting as highly mineralized inclusions (Fig. 3C, arrows). Additionally, vertebrae in 1-year-old *chd7*^-/-^ zebrafish presented with a larger bone volume fraction (BV/TV) compared to controls (Fig. 3G-I). Analysis of the vertebrae neural arch revealed distortion with a significantly larger rising angle, as well as a larger variance of the angles (Fig. 3F). Vertebrae showed a significantly changed body angle and significant variance of body angle and vertebrae length (Supplemental Table 3). While a depletion of vBMD in the arches and abnormal vertebrae body structure was observed in 2-year-old fish, the effects seen on total vBMD and BV were not apparent in the few samples that survived until this stage (Supplemental Fig. 2 and Supplemental Table 4). Notably, one of our *chd7*^-/-^ mutants showed vertebral fusion of the analyzed precaudal vertebra, which was not observed in any of the controls (Supplemental Fig. 1). This further demonstrates the wide variation in severity of the bone phenotype in *chd7*^-/-^ zebrafish.

**Fig 3.**
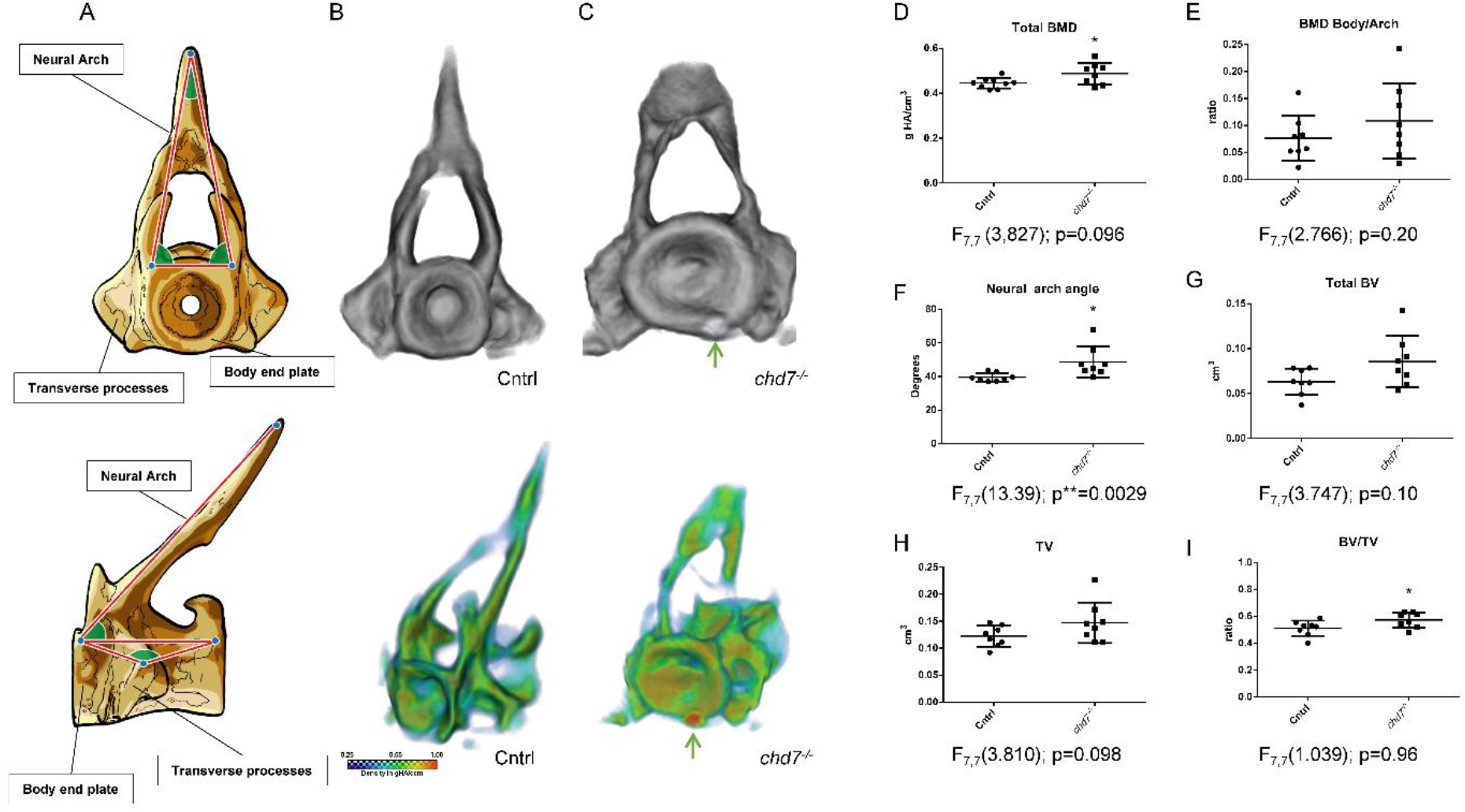
Precaudal vertebrae show inefficiency for proper mineralization. A. Sketch diagram of precaudal vertebrae (frontal and lateral) indicating structure and measured angles. B-C. (top) Individual rendering of precaudal vertebrae, frontal view of tissue over 0.41 g HA/cm^3^ threshold and (bottom) vBMD intensity map (Range blue-red; 0.41 – 1.00 g HA/cm^3^) showing morphological abnormalities and growth zone malformations with highly mineralized inclusions (arrow) D. vBMD of whole vertebrae showing increasing density with increasing age and e. increasing ratio of vBMD arch/vertebrae body. F-H neural arch angle, BV, TV and ratio BV/TV. (n=8/genotype were analyzed). Significance of student t-test with Welch’s correction is included in the graphs with: *p<0.05; **p<0.01. F-test results are indicated underneath as F_degree of freedom numerator, degree of freedom denominator_ (F-value); p-value: *p<0.05; **p<0.01; ***p<0.001

### Abnormal caudal vertebrae in chd7^-/-^ zebrafish vary in arch angles and reduced mineralization

We next determined key structural features (Fig. 4A) and mineralization characteristics of caudal vertebrae. The *chd7*^-/-^ zebrafish presented with warped hemal and neural arches and abnormal body structure compared to controls (Fig. 4B, C). Similar to precaudal vertebrae, the growth zones of the body end plates were enlarged and showed inclusions of highly mineralized matrix (Fig. 4C, arrows). Unlike control zebrafish, *chd7*^-/-^ mutants had abnormal vertebrae body structure, which usually exhibited an hourglass shape and resulting in greater variance of measured body angles (Supplemental Table 3).

**Fig 4.**
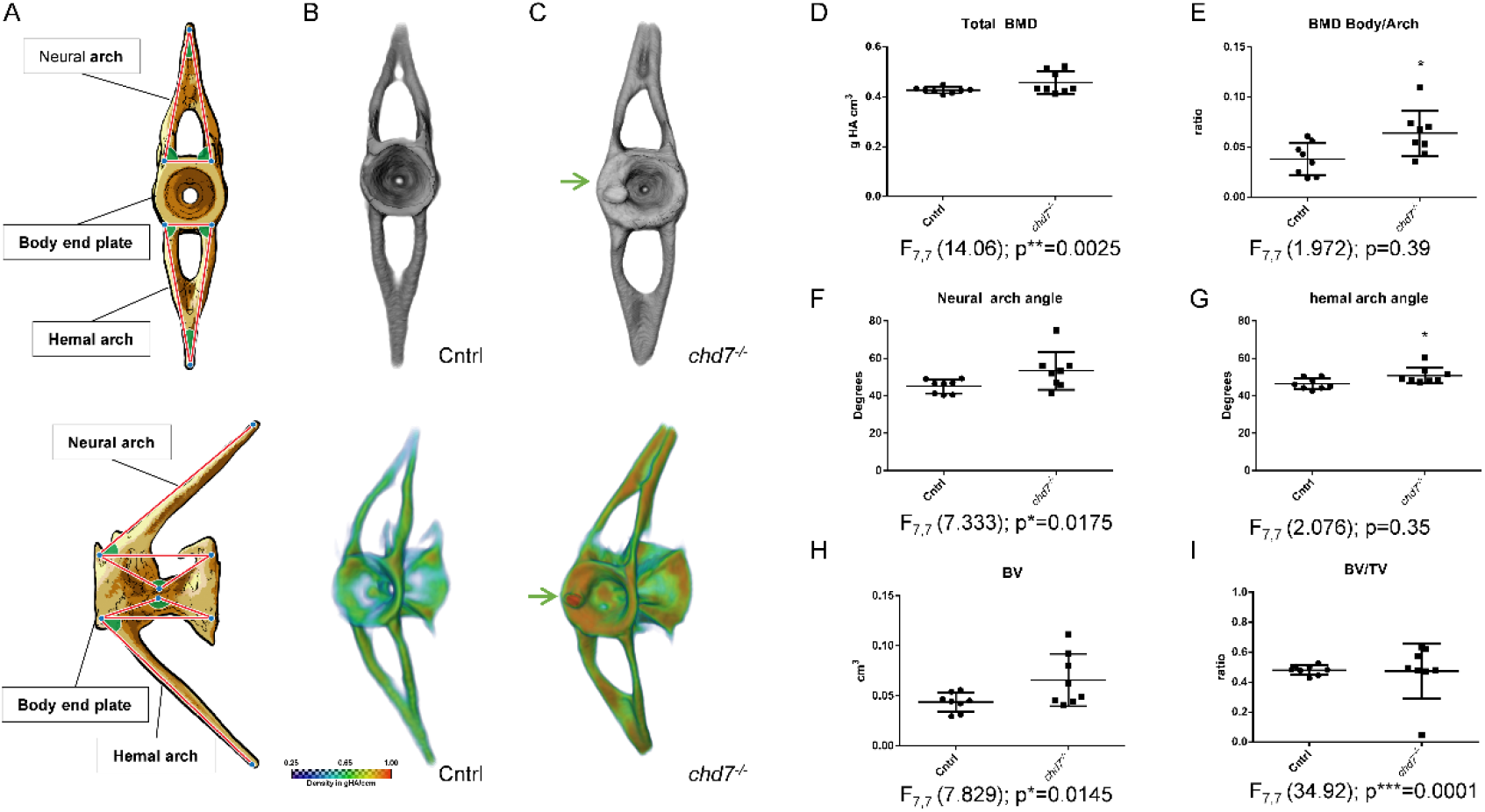
Caudal vertebrae show abnormal structure. A. Sketch diagram of precaudal vertebrae indicating structure and measured angles (frontal and lateral). B-C. (top) Individual rendering of caudal vertebrae, frontal view of tissue over 0.41 g HA/cm^3^ threshold and (bottom) vBMD intensity map (Range blue-red; 0.41 – 1.00 g HA/cm3) showing morphological abnormalities and growth zone malformations with highly mineralized inclusions (arrow) D. vBMD of whole vertebrae and E. increasing ratio of vBMD arch/vertebrae body. F-H neural arch angle, BV, TV and ratio BV/TV. (n=8/genotype were analyzed). Significance of student t-test with Welch’s correction is included in the graphs with: *p<0.05; **p<0.01. F-test results are indicated underneath as F_degree of freedom numerator, degree of freedom denominator_ (F-value); p-value: *p<0.05; **p<0.01; ***p<0.001

In line with the neural arch screened in the precaudal vertebrae, the hemal arch in the caudal vertebrae showed significantly wider arch rising angles (Fig. 4F), while the neural arch displayed significantly greater variance (Fig. 4G) in 1-year-old *chd7*^-/-^ zebrafish. In contrast to precaudal vertebrae data, the caudal vertebrae showed significantly larger variance of Total vBMD and of both arches and vertebral body vBMD (Supplemental Table 3). A significant insufficiency in mineralization of the arches compared to the vertebrae body was also measured in *chd7*^-/-^ zebrafish compared to controls (Fig. 4D, E). Further analysis also revealed a significant variance in BV, TV and BV/TV in *chd7*^-/-^ zebrafish compared to controls (Fig. 4H, I).

While significantly distorted arches were still observed in 2-year old mutants, the effects on vBMD and BV were not noticed in the few surviving fish that could be screened (Supplemental Fig. 3 and Supplemental Table 4).

### Col2a1 of the ECM is depleted in chd7^-/-^ zebrafish

To perform a more complete analysis of the skeletal phenotype, we also examined the extracellular matrix (ECM) of bone structures in 6 dpf larvae and 1-year-old *chd7*^-/-^ zebrafish. Gene expression analysis at 9 dpf revealed a significant downregulation of the ECM gene *col2a1a* (Fig. 5E). Antibody staining showed that all craniofacial structures are already depleted of the key Col2a1a protein by 5 dpf (Fig. 5A). Vacuole formation within the structures was apparently unimpaired. In adult *chd7*^-/-^ zebrafish, immunofluorescence showed a striking depletion of Col2a1 in the cartilage of vertebrae seen in the Weberian and precaudal vertebrae, with almost abolished Col2a1 signal (Fig 5B). This pattern was effectively reproduced in all regions of the spine of *chd7*^-/-^ zebrafish.

**Fig 5.**
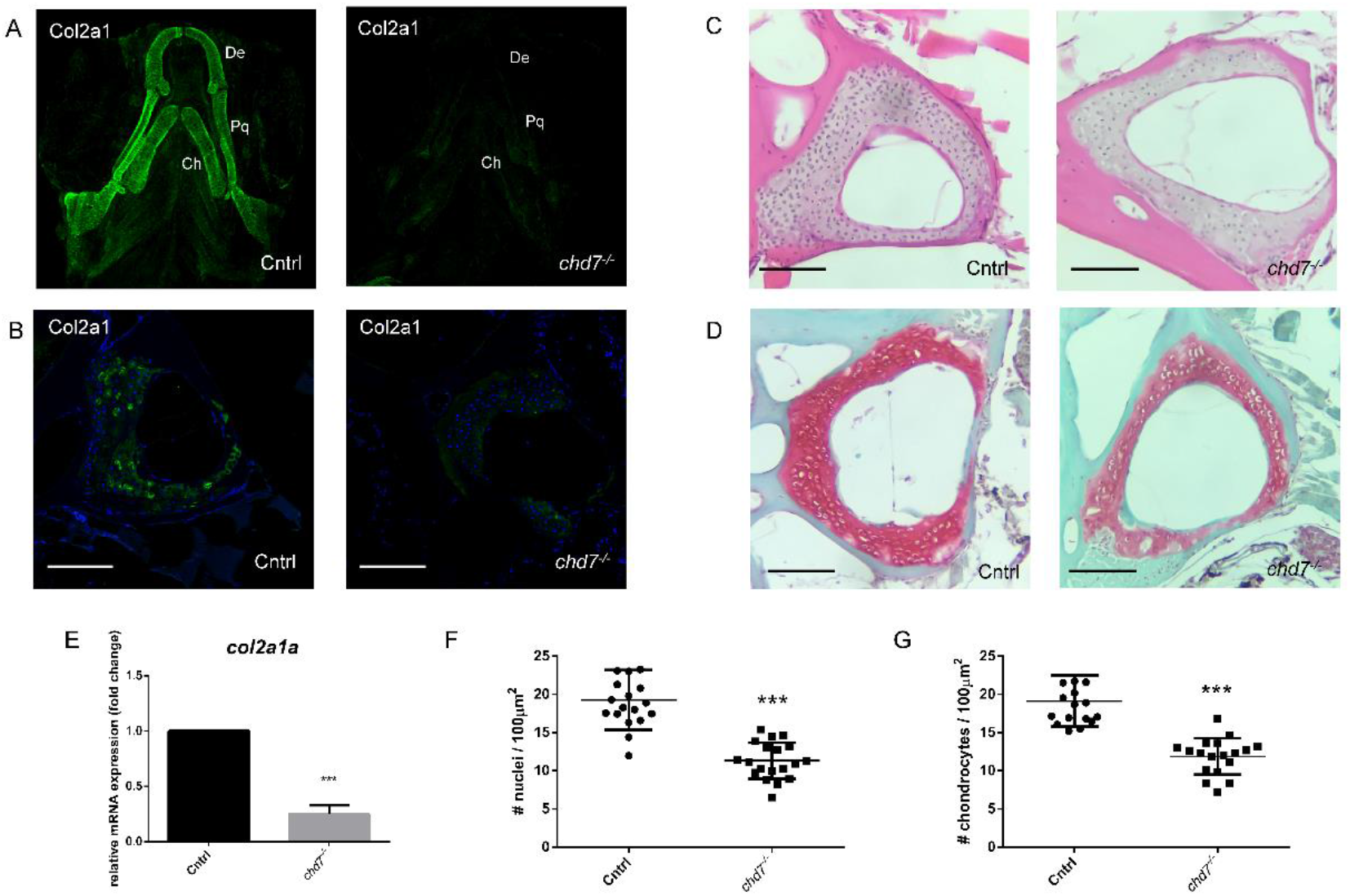
ECM Collagen2a1 deficiency in larvae and 1-year-old *chd7*^-/-^ adults. A. Immunofluorescence of Col2a1 in 5 dpf Cntrl (n=9) and *chd7*^-/-^ (n=8) larvae, ventral view of craniofacial cartilage (De: dentary; Pq: palatoquadrate; Ch: ceratohyal) B. Immunofluorescence of Col2a1 in precaudal vertebrae sections of Cntrl and *chd7*^-/-^ mutants with Col2a1 in green and DAPI in blue. C. H&E staining of precaudal vertebrae section of Cntrl and *chd7*^-/-^ mutants D. Safranin O/fast green staining of Weberian and precaudal vertebrae sections of Cntrl and *chd7*^-/-^ mutants showing cartilage in red. E. RT-qPCR of *col2a1a* at 9dpf. F. Total nuclei count in Weberian and precaudal vertebral cartilage detected in H&E staining per 100μm^2^ in Cntrl (N=4, n=17) and *chd7*^-/-^ (N=4, n=18) G. Chondrocyte nuclei count in precaudal vertebral cartilage detected in Safranin O/fast green staining per 100μm^2^ Cntrl (N=4, n=18) and *chd7*^-/-^ (N=4, n=19) Scale bar presents 100μm. Significance of student t-test with: *p<0.05; **p<0.01 ***p<0.001

We tested further for cell nuclei and chondrocytes in vertebral cartilage. Hematoxylin-Eosin (HE) staining of Weberian apparatus and precaudal vertebrae sections showed a significant reduction in the number of nuclei measured in the vertebral cartilage of 1-year-old *chd7*^-/-^ zebrafish and controls (Fig. 5C, F). Complementary analysis using Safranin/Fast Green O staining, revealed a significant reduction in the number of chondrocytes in the vertebral cartilage in *chd7*^-/-^ zebrafish in line with the HE staining (Fig. 5D, G). Altogether, the findings indicate an altered cartilage composition in addition to the observed bone deformations.

## Discussion

### Efficiency of model

CS is a multisystemic disorder likely involving a variety of underlying mechanisms that include CHD7. Therefore, different preclinical models are required to individually identify the various molecular mechanisms. Even though mouse models for CS exist, none have been reported to show the bone phenotype observed in CS patients. Our study presents a *chd7* deficient zebrafish that has a bone phenotype resembling that of CS patients, which we used to investigate skeletal development in CS. CS patients commonly present with square face, broad nasal bridge, small mouth and facial asymmetry (9). While scoliosis in CS patients has been attributed to poor muscular development, our study proposes an additional factor involving impaired early mineralization of bone in the vertebrae. In line with clinical features exhibited by CS patients, our mutants display craniofacial dysmorphism with reduced mineralization and spinal deformities indicative of scoliosis (10–12). Interestingly, our analysis revealed that craniofacial and vertebral abnormalities are already present in young larvae of *chd7*^-/-^ zebrafish. Additional clinical studies will be required to investigate the onset of skeletal deformities in CS patients. However, early analysis of bone density and spinal development has been suggested by CS guidelines (45, 46). This early and detailed analysis may reveal less severe or early onset phenotypes. Similarly, we observed that adult *chd7*^-/-^ zebrafish, which appeared without spinal deformities on visual inspection, showed signs of spinal abnormalities in microCT analysis, suggesting that mostly mild phenotypes reach this age. This is supported by the high lethality rate shown by Jamadagni et al. (unpublished), yet the underlying cause of lethality is still unclear. Even though 5 dpf old larvae were significantly smaller than their control counterparts, this is expected as part of the CHARGE phenotype. Nonetheless, overall morphology suggests that they are not in a notable developmental delay. Unfortunately, key “landmarks” for development as previously proposed are expectedly affected in *chd7*^-/-^ zebrafish, including overall growth, pigment migration, swimbladder inflation and mineralization (47). However, we can conclude if the mineralization effect observed at 9dpf was in fact developmental delay, we would expect mineralization to recover. Yet, the phenotype was still evident in later development (Supplemental Fig 1. D).

Progressive skeletal disorders such as osteopenia and osteoporosis are characterized by a reduction in bone mineralization over time leading to increased fracture risk (48). CS patients have been shown to present with significantly reduced bone mineral density (13). Correspondingly, reduced vertebrae mineralization in *chd7*^-/-^ zebrafish by 12 dpf was observed in a previous study involving morpholinos in zebrafish which had observed a reduced mineralization of vertebral structures (27). Confirming previous data, we observed insufficient mineralization of vertebra along the posterior spinal column by 4 weeks of age. In earlier analysis of larvae at 6 dpf, we were not able to detect a delay in the onset of mineralization, but already by 9 dpf we saw a significant decrease in the number of calcified vertebrae. Furthermore, mineralization appeared to be less efficient in posterior vertebrae, which could signify an impairment in ossification, since vertebrae in zebrafish undergo ossification from an anterior to posterior direction (49). An exception from this rule is the Weberian vertebrae C1 and C2, which are ossified later in wildtypes but earlier in *chd7*^-/-^ mutants. The craniofacial abnormalities we observed are in line with known neural crest deficiencies in CS which are known to affect cartilage and bone development (50).

### Nutrient dependency

Our study showed that the onset of calcification was not delayed in *chd7* deficient zebrafish. However, progressing calcification was decreased already 4 days post feeding onset, even though access to food was identical in control and mutants. In contrast, differences in calcification of vertebrae in control and mutants without access to food was not significant. Previous studies have shown that feeding delay until 9 dpf does not affect future fish growth and viability and was therefore our cut-off point(51). This data suggests that possibly an underlying mechanism involving either reduced uptake or metabolizing nutrients may play a part in the observed phenotype and this link remains to be investigated. Bone calcification and mineralization is dependent on proper nutrition and is often discussed regarding optimal bone health in humans (52). Insufficient levels of calcium and nutritional uptake are considered an increased risk for bone injury in CS patients (11). To optimize nutritional uptake, the supplementation of calcium and other nutrients to counteract poor bone health has been proposed for CS patients (53). In fact, case reports show supplementation with calcium and vitamin D significantly increased aBMD in CS patients (54). A possible nutrient deficiency in CS, could be linked to gastrointestinal nutrient uptake difficulties in *chd7*^-/-^ zebrafish (55), yet this needs to be further investigated. Thus, this possible deficiency in nutrient uptake could further hinder ossification in *chd7*^-/-^ mutants. We propose using our zebrafish model of CS model to test the effect of guidelines such as calcium and nutrient supplementation, nutrient metabolism and mechanical loading in form of swim training to enhance bone strength in CS.

### Loss of chd7 reduces osteogenesis in zebrafish

Osteoblast precursors commonly express differentiation markers such as *runx2* and *Osterix/sp7* (56). Upon maturation of these osteoblasts, expression of mineralization related genes such as *bglap* (osteocalcin) are expressed. Analysis of these three genes showed that they were all significantly downregulated in *chd7*^-/-^ larvae. This suggests a deficiency in mature osteoblasts, required for the mineralization of facial bone structures and early spinal column. Misregulation or loss of these genes, such as *osterix/sp7* has been directly linked to poor bone mineralization in zebrafish and humans (57–59). Notably, among all osteogenesis related gene expressions significantly reduced in *chd7*^-/-^ larvae, *sost*, *ctsk, acp5* and *col2a1* have been identified as direct regulatory targets of Chd7 by Chip-Seq (60). Furthermore, the regulation of osteogenesis has been linked to a regulatory complex with PPARγ (23). Cell culture experiments have proposed the role of CHD7 in osteogenesis by inhibiting PPARγ target genes by a protein complex involving CHD7, NLK and SETDB1. Upon loss of CHD7 this complex will fail to inhibit PPARγ target genes and therefore inhibit osteogenesis in preference over adipogenesis (23). Strikingly, our qPCR results are consistent with downregulation of Pparγ target genes such as *runx2a* and *runx2b*. Furthermore, gene expression relevant for bone calcification and remodelling are misregulated such as *sost* and *ctsk*. Thus, our study is the first to indicate the conservation of the proposed pathway shown in cell culture in an *in vivo* model.

### Weberian apparatus

The Weberian apparatus is a specialized structure that connects the auditory system and the swim bladder in a complex with the first 4 vertebrae to amplify sound vibrations (33). Our results show that key structures in the apparatus fail to efficiently form and/or mineralize. Two of these structures, the tripus and the parapophysis, are formed to connect to the swim bladder. Both are malformed in all tested *chd7*^-/-^ mutants to varying degrees. This malformation may imply an inefficiency in its function concerning both the swim bladder and auditory system. Ear abnormalities are a major symptom in CS patients and functionality is affected. Furthermore, the *chd7*^-/-^ mutant shows swim bladder defects (Breuer et al., unpublished) that could be additionally affecting the morphology in tripus and parapophysis or vice versa. Even though the Weberian apparatus is extremely specialized for carp and carp-like fish (Ostariophysi), it is likely that it underlies the same pathways for osteogenesis as other parts of the spine.

### Abnormal vertebrae and vBMD variability

We examined volumetric bone mineral density (vBMD) and other parameters (e.g. BV, TV) in vertebral bone structures. Most strikingly, we detected large variance in density and all parameters examined, as well as some significant changes in both precaudal and caudal vertebrae. The large variance observed in *chd7*^-/-^ mutants is in line with CS patients showing varying severity in spinal deformities (12, 61). Furthermore, previous studies have shown that even minor changes in the described parameters can impact the risk for spinal deformities and fractures in disease and under treatment (62, 63).

Another interesting feature we observed in most *chd7*^-/-^ mutants, were highly mineralized inclusions, most commonly, localized towards the growth zone of the vertebrae. We did not identify the source of these highly mineralized structures, but they could be caused by localized clusters of osteoblasts during vertebrae mineralization. Another possibility is the occurrence of calcified cartilage, which is more mineralized in comparison to bone. However, this is highly speculative and histologic analysis of such small and highly localized structures will be challenging. Since these inclusions are highly mineralized, it seems likely that their appearance impairs the bone structurally as higher mineralized collagen is stiffer, yet less tough (64). It will be of interest to see if future studies can take a closer look at these previously unidentified features. While these minor defects could be involved in the structural integrity of the spine in CS patients, these minor features may be missed during dual-energy x-ray absorptiometry (DEXA) screens of CS patients when assessing aBMD.

Analysis revealed a decreased mineralization in the caudal vertebral arches of the mutant fish. This likely reduces the mechanical competence of these vertebrae in comparison to healthy vertebrae. A compromised mechanical competence alongside low BMD also increases fracture risk and can be an underlying cause of scoliosis (65–67). Unfortunately, natural history BMD data of CS patients throughout early development is not available, to the best of our knowledge. However, aBMD analysis of patients with idiopathic scoliosis in CS via DEXA scans has been studied and shows significant reduction in some cases (13, 68). In general, there is an increased fracture risk and incidence of scoliosis corresponding to decreased BMD, such as in osteopenia and osteoporosis (48, 66). In line with this reduced mineralization an increased fracture and injury risk has been reported for CS patients (13). Accordingly, proper observation and focus on early DEXA analysis is considered for CS guidelines (45, 46). Our microCT analysis of zebrafish vertebra is not without limitations. An isotropic voxel size of 10.6 μm cannot resolve cancellous bone structures thus overestimating bone volume. However, this limitation applies systemically so differences between groups hold true. Other outcomes such as BMD and shape are less affected by resolution.

Our analysis throughout development of *chd7*^-/-^ deficient zebrafish suggests that only mild phenotypes survive onto later stages with severity decreasing as age progresses. However, the high lethality in both patients and our model is connected to a multisystemic phenotype in which one mechanism is unlikely to be the exclusive underlying cause for this characteristic. The link between Chd7 and severe spinal and vertebrae abnormalities, including kyphosis has been observed in our samples (7, 9).

### Collagen and chondrocyte depletion as novel risk factor

Our data in *chd7* deficient zebrafish indicate that reduced vBMD may not be the only underlying risk factor for idiopathic scoliosis in CS patients. Furthermore, craniofacial development in CS may be dependent on ECM matrix composition. Bone matrix integrity may be weakened by a decrease in the collagen component Col2a1 in the craniofacial and vertebral cartilage of *chd7* deficient zebrafish. Col2a1 is required in the ECM of various cell types including osteoblasts, chondrocytes, external ligament connective tissue cells, as well as notochord basal cells and expressed in the corresponding tissues (69, 70). Each cell type is required for proper bone formation and could likely contribute to spinal injuries and malformations. Furthermore, *col2a1a* expression is observed throughout ossification of the notochord (71). Reduction in this collagen matrix in the notochord can affect proper calcification and is known to cause spondyloepiphyseal dysplasia congenita, which affects skeletal and spine development (72). Strikingly, *sedc* mice, which are deficient in *Col2a1*, show enlargement and malformations in the growth plates of vertebrae and degeneration of cartilage, similar to our study in zebrafish (73). *Col2a1*a is expressed in osteoblasts and chondrocytes of teleosts and regulated by *sox9*, which in turn is known to be regulated by *chd7* (71, 74–76). Sox9 is also well known to be required in craniofacial development, such as its related factor Sox10, both of which have been connected to the craniofacial phenotype in CHARGE (77). The hypertrophic morphology of the chondrocytes seen in our study may further indicate signs of an altered matrix composition and may contribute to an increased risk in osteoporosis (78, 79).

In summary, our study is the first to identify skeletal abnormalities linked to *chd7* deficiency. These abnormalities include morphological changes in the skull and vertebrae, reduced bone mineralization, impaired osteoblast activity, highly mineralized inclusions and Col2a1 deficiency in the *chd7*^-/-^ zebrafish model organism. The use of this preclinical model will allow for future studies to further understand the underlying mechanisms of spinal deformities in CS.

## Materials and Methods

### Zebrafish husbandry

Wildtype (Cntrl) and mutant (*chd7*^-/-^) zebrafish were kept at 28°C in a 12h/12h dark/light cycle and maintained in accordance with Westerfield et al.(80). All zebrafish in this study were fed a steady diet of Skretting^®^GemmaMicro starting at 5 dpf. Embryos were raised at 28.5°C and staged as previously described by Kimmel et al.(81) All experiments were performed in with the guidelines of the Canadian Council for Animal Care and the local ethics committee.

### Skeletal stainings and analysis of larvae

Calcein staining was done as described(38). In short, zebrafish larvae at 6 or 9 dpf were exposed to calcein (2g/L) (Sigma-Aldrich) in water for 10 minutes. Larvae were then washed at least 3 times in fish water for 10 minutes to remove excess calcein. Larvae were then anesthetized using tricaine before being imaged using an AxioZoom V16 (Zeiss). For nutrition dependency tests, larvae were either deprived of food until 9 dpf or fed a regular Skretting^®^GemmaMicro diet starting at 5 dpf.

Alcein blue staining was performed according to (82).

Alizarin Red was performed as previously shown (83). In brief, larvae were fixed for 2 hours with 2% PFA. The PFA was then removed by washes in PBS 3×10 minutes and then placed in 25% glycerol/0.1%KOH. Finally, larvae were stained with 0,05% Alizarin Red (Sigma-Aldrich) in H2O for 30 minutes and stored in 50% glycerol in 0.1%KOH.

### MicroCT Imaging of the spinal and craniofacial skeleton

Adult zebrafish were euthanized using tricaine (MS-222, Sigma-Aldrich). Tissue was fixed in 4% PFA prior to microCT analysis. Ex vivo microCT at an isotropic voxel size of 10.6 μm (SkyScan1276, Bruker, Kontich, Belgium) was performed to assess differences in Weberian and vertebral bone mass, mineral density, and microstructure (55 kVp, Al 0.25 mm filter, 200 μA source current, 0.3° steps for full 360°). Additionally, ex vivo microCT of craniofacial bone structures at an isotropic voxel size of 5.0 μm was performed (55 kVp, Al 0.25 mm filter, 72 μA source current, 0.16° steps for full 360°) to assess morphological features of the skull… Images were reconstructed using standard reconstruction algorithms provided with the microCT. Spinal and craniofacial deformities of controls and mutants were analyzed of 1-year (n=8 for vertebrae, n=5 for skull) and 2-year-old (n=5) zebrafish.

### Analysis and bone density/volume/angles

From the reconstructed microCT images of the whole spine, the Weberian apparatus, as well as vertebrae 7 (abdominal) and 20 (caudal) were segmented. Further, the intercalarium, the tripus as well as the parapophysis were segmented from the Weberian apparatus. For the vertebrae, the neural as well as the hemal arch were segmented from the vertebral body. To differentiate between bone and background, a density-based threshold of 0.41 gHA/cm^3^ determined by Otsu’s method(84) was used. For each segmented bone in the Weberian apparatus outcomes included: bone volume BV (mm^3^), tissue volume TV (mm^3^), bone volume fraction BV/TV (mm^3^/mm^3^) and volumetric bone mineral density vBMD (gHA/cm^3^). For abdominal and caudal vertebrae the outcomes included: bone volume BV, tissue volume TV, bone volume fraction BV/TV, total bone mineral density T.vBMD, vertebral body vBMD VB.BMD, vertebral arch vBMD VA.BMD, as well as arch-body angle, body angle, arch opening angle and rising angles (see Fig.2A and 3A). For caudal vertebrae, the angle outcomes were calculated for hemal as well as neural arches. Craniofacial outcomes included: mandibular arch length, complete skull length measured from the tip of the dentary to the furthest point of the supraoccipital, width of the skull, measured between the opercular tips as well as mandibular arch angle, and craniofacial angle.

### qRT-PCR

Total RNA was isolated from 9 dpf old zebrafish larvae using TriReagent. 1μg of RNA was used for cDNA synthesis using cDNA vilo kit. qRT-PCR was performed with SYBR Green mix (BIORAD) with a Lightcycler96^®^ (Roche). Gene expression was analyzed relative to the housekeeping gene elf1α. Primers used for *runx2a, runx2b, sost, bglap, sp7/osterix, mgp, postna, acp5a, ctsk, col2a1 and mgp* are shown in supplementals Table 5.

### Spinal sections/Stainings/Immunohistochemistry

The zebrafish were euthanized in tricaine (MS-222, Sigma-Aldrich) and fixed for 3 days in 4% paraformaldehyde at 4°C, after the abdomen was opened to ensure proper fixation. The spinal cords were dissected and decalcified with EDTA 10 % for 3 days under agitation at RT. After fixation and decalcification, the zebrafish were embedded in paraffin. Longitudinal sections (5μm) were obtained and were deparaffinized in xylene and were rehydrated in a graded series of ethanol. The slides were stained with hematoxylin (STATLAB Medical Products, LLC) for 4 min, and washed with alcohol-acid, and were rinsed with tap water. In the blueing step, the slides were soaked in saturated lithium carbonate solution for 10 sec, and then rinsed with tap water. Finally, staining was performed with eosin Y (STATLAB Medical Products, LLC) for 2 min and mounted with Permount™ mounting medium.

For Safranin O/Fast green, after dehydration, the slides were stained with Weigert’s hematoxylin (Sigma-Aldrich) for 4 min, wash under tap water, immersed in alcohol–acid for 5 sec, then washed under water. The slides were stained with Fast green 0.02 % for 2 min, washed with acetic water 1% for 20 sec, then stained directly in safranin 0.01 % for 5 min and mounted with Permount™ mounting medium. Images were taken using an AxioZoom V16 (Zeiss) and area and number of nuclei were determined using ImageJ.

Immunofluorescence for Col2a1 was performed as previously shown. For 5 dpf larvae fish were fixed with 4% PFA overnight. Samples were blocked with 4% BSA in PBT (1% Triton) and samples placed in primary antibody for Col2a1 (1 in 20; Developmental studies hybridoma bank) at 4°C overnight. Then washed with PBT for several hours and then incubated with secondary antibody Goat-anti-Mouse AlexaFluor 488 (1 in 200) at 4°C overnight.

For 1-year old paraffin sections, epitope demasking was performed by digestion with Proteinase K (20μg/ml). Tissue was blocked with NGS and samples treated with primary antibody for Col2a1 (1 in 10; Developmental studies hybridoma bank) and secondary antibody Goat-anti-Mouse Alexafluor 488 (1 in 300). Counter stain was done using DAPI-mounting medium.

Images were taken using a LSM780 (Zeiss).

### Statistics

Statistical analysis was carried out using Prism-GraphPad^®^ (GraphPad, San Diego, CA, USA). qPCR data and nuclei quantification were tested with student’s t-test with Welch’s correction. Calcification analysis in larvae was performed with one-way ANOVA and post-hoc Tukey Test. MiroCT data of Control vs *chd7*^-/-^ mutants was analyzed using student’s t-test with Welch’s correction and Variance determined by F-test analysis. Significance was determined at *<0.05; **<0.01 and ***<0.001

## Acknowledgements

The authors would like to thank Beatrice Steyn and Priyanka Jamadagni for experimental support, and Mark Lepik for graphic design. SP is supported by the Natural Science and Engineering Research Council (NSERC), Canadian Institutes of Health Research (CIHR), an ALS Canada-Brain Canada Career Transition Award and a FRQS Junior 1 research scholar. BMW is supported by FRQS Programme de bourses de chercheur. MR is supported by FRQS Programme de bourses de chercheur. We also thank Shriners Hospital for Children and the CIHR (165939) for financial support.

## Author roles

MB, MR, BMW and SP designed the study. MB, MR and CZ performed the experiments. MB and MR drafted the manuscript. MB, MR, CZ, BMW, SP reviewed and edited the manuscript.

