## Supplemental Data for "Abnormal craniofacial and spinal bone development with *col2a1*a depletion in a zebrafish model of CHARGE syndrome"

### Supplemental Figures

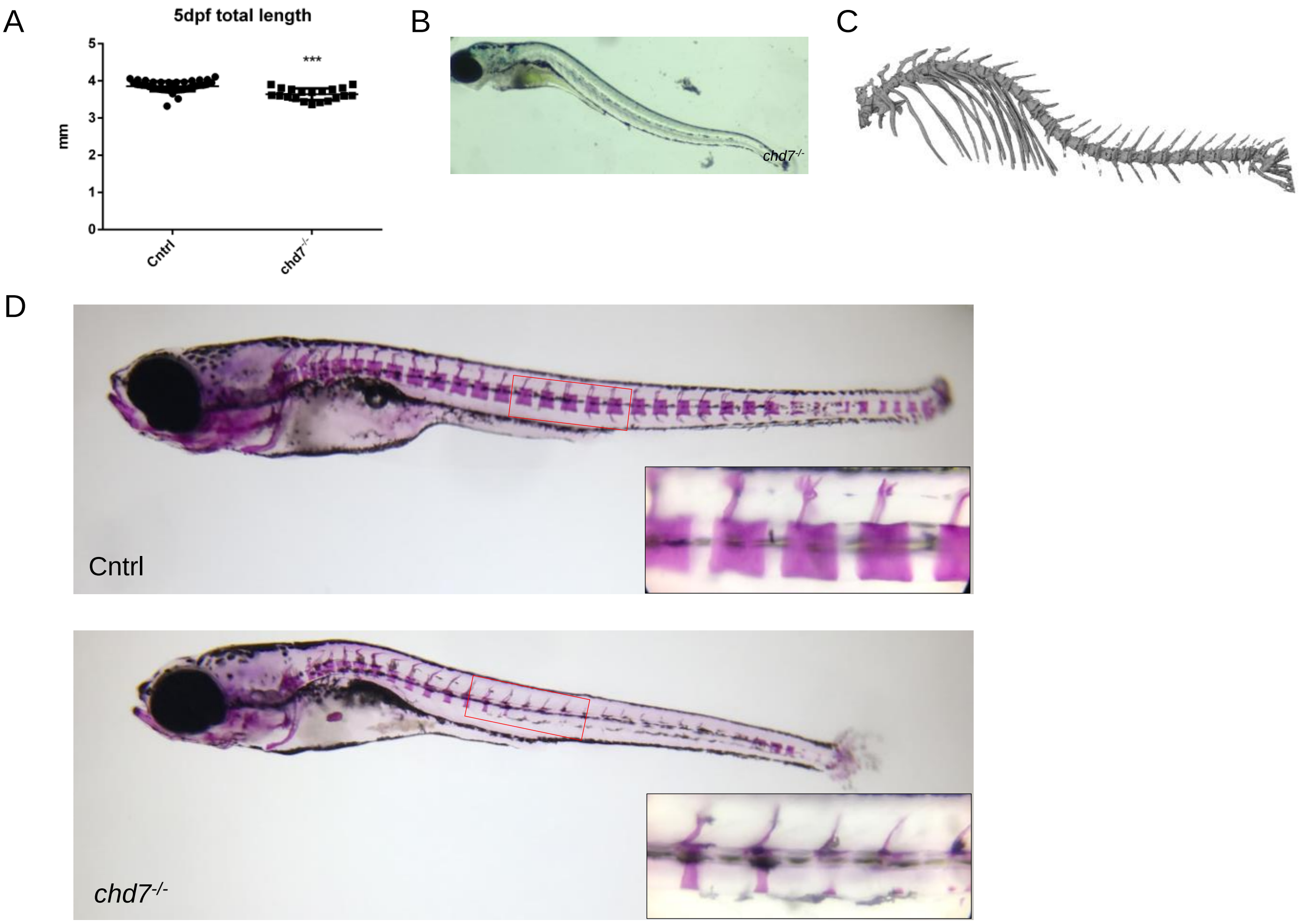


Supplemental Fig 1. Spinal analysis in *chd7* mutants A. Total length of 5dpf larvae B Lateral image of *chd7* larvae presenting with severe kyphosis at 9 dpf. C. Severe scoliosis in an adult *chd7* zebrafish D. Alizarin Red staining of 1 month old Cntrl and *chd7* zebrafish with close up of indicated ROI

#### Supplemental Figures

Supplemental Table 1. Statistical analysis of mineralization of the Weberian apparatus in 1-year-old zebrafish

| | cntrl 1-year-old | | | CHD7$^{-/-}$ 1-year-old | | | | |
|---|---|---|---|---|---|---|---|---|
| *Weberian structures* | n = 8 | | | n = 8 | | | *p(t-test)* | *p(F-test)* |
| **Intercalarium.TV [mm$^3$]** | 0.046 | ± | 0.009 | 0.056 | ± | 0.020 | 0.210 | 0.048 |
| **Intercalarium.BV [mm$^3$]** | 0.028 | ± | 0.007 | 0.035 | ± | 0.014 | 0.261 | 0.096 |
| **Tripus.TV [mm$^3$]** | 0.085 | ± | 0.025 | 0.085 | ± | 0.048 | 0.987 | 0.101 |
| **Tripus.BV [mm$^3$]** | 0.051 | ± | 0.024 | 0.047 | ± | 0.038 | 0.823 | 0.244 |
| **Parapophysis.TV [mm$^3$]** | 0.199 | ± | 0.037 | 0.213 | ± | 0.100 | 0.710 | 0.018 |
| **Parapophysis.BV [mm$^3$]** | 0.136 | ± | 0.036 | 0.145 | ± | 0.081 | 0.777 | 0.048 |

Supplemental Table 2. Statistical analysis of mineralization of the Weberian apparatus in 2-year-old old zebrafish

| | cntrl 2-year-old | | | CHD7$^{-/-}$ 2-year-old | | | |
|---|---|---|---|---|---|---|---|
| *Weberian structures* | n = 5 | | | n = 5 | | | *p(t-test)* |
| **Intercalarium.BMD [gHA/cm$^3$]** | 0.531 | ± | 0.031 | 0.531 | ± | 0.024 | 0.979 |
| **Intercalarium.TV [mm$^3$]** | 0.076 | ± | 0.032 | 0.063 | ± | 0.008 | 0.416 |
| **Intercalarium.BV [mm$^3$]** | 0.049 | ± | 0.022 | 0.040 | ± | 0.006 | 0.445 |
| **Intercalarium.BV/TV** | 0.637 | ± | 0.032 | 0.641 | ± | 0.028 | 0.837 |
| **Tripus.BMD [gHA/cm$^3$]** | 0.498 | ± | 0.043 | 0.486 | ± | 0.069 | 0.751 |
| **Tripus.TV [mm$^3$]** | 0.115 | ± | 0.046 | 0.113 | ± | 0.047 | 0.963 |
| **Tripus.BV [mm$^3$]** | 0.073 | ± | 0.036 | 0.070 | ± | 0.039 | 0.885 |
| **Tripus.BV/TV** | 0.624 | ± | 0.048 | 0.592 | ± | 0.075 | 0.449 |
| **Parapophysis.BMD [gHA/cm$^3$]** | 0.592 | ± | 0.036 | 0.590 | ± | 0.014 | 0.949 |
| **Parapophysis.TV [mm$^3$]** | 0.334 | ± | 0.185 | 0.220 | ± | 0.028 | 0.242 |
| **Parapophysis.BV [mm$^3$]** | 0.242 | ± | 0.151 | 0.153 | ± | 0.020 | 0.262 |
| **Parapophysis.BV/TV** | 0.706 | ± | 0.042 | 0.695 | ± | 0.006 | 0.565 |

### Supplemental Figures

Supplemental Table 3. Statistical analysis of mineralization of the precaudal and caudal vertebrae in 1-year-old zebrafish

| c | cntrl 1-year-old | | | CHD7$^{-/-}$ 1-year-old | | | | |
|---|---|---|---|---|---|---|---|---|
| *precaudal vertebra* | n = 8 | | | n = 8 | | | *p(t-test)* | *p(F-test)* |
| **Arch.vBMD [gHA/cm$^3$]** | 0.369 | ± | 0.043 | 0.356 | ± | 0.047 | 0.593 | 0.995 |
| **Body.vBMD [gHA/cm$^3$]** | 0.445 | ± | 0.050 | 0.464 | ± | 0.064 | 0.520 | 0.643 |
| **ArchOpen.angle [°]** | 20.4 | ± | 2.5 | 26.2 | ± | 10.6 | 0.171 | 0.002 |
| **ArchRise1.angle [°]** | 80.0 | ± | 3.0 | 76.7 | ± | 6.0 | 0.195 | 0.128 |
| **ArchRise2.angle [°]** | 80.4 | ± | 2.0 | 77.4 | ± | 5.7 | 0.198 | 0.019 |
| **Diff.R1-R2 [°]** | 3.9 | ± | 2.5 | 4.8 | ± | 3.9 | 0.618 | 0.361 |
| **Body.angle [°]** | 134.5 | ± | 5.1 | 133.8 | ± | 11.1 | 0.869 | 0.081 |
| **Vertebra.length [µm]** | 0.601 | ± | 0.230 | 0.670 | ± | 0.084 | 0.472 | 0.011 |
| *caudal vertebra* | n = 8 | | | n = 8 | | | *p(t-test)* | *p(F-test)* |
| **Arch.vBMD [gHA/cm$^3$]** | 0.413 | ± | 0.020 | 0.433 | ± | 0.052 | 0.334 | 0.027 |
| **Body.vBMD [gHA/cm$^3$]** | 0.451 | ± | 0.022 | 0.486 | ± | 0.055 | 0.129 | 0.040 |
| **NeuralBody.angle [°]** | 133.7 | ± | 4.2 | 133.9 | ± | 11.5 | 0.965 | 0.023 |
| **NeuralArchOpen.angle [°]** | 17.4 | ± | 1.7 | 19.1 | ± | 3.5 | 0.265 | 0.091 |
| **NeuralArchRise1.angle [°]** | 82.2 | ± | 2.9 | 79.9 | ± | 3.0 | 0.146 | 0.955 |
| **NeuralArchRise2.angle [°]** | 80.7 | ± | 2.5 | 81.5 | ± | 2.5 | 0.547 | 0.835 |
| **Diff.Neural.R1-R2 [°]** | 4.1 | ± | 3.0 | 3.7 | ± | 2.5 | 0.802 | 0.523 |
| **HemalBody.angle [°]** | 127.0 | ± | 4.2 | 127.7 | ± | 2.0 | 0.684 | 0.061 |
| **HemanArchOpen.angle [°]** | 20.3 | ± | 2.2 | 24.2 | ± | 4.9 | 0.071 | 0.070 |
| **HemalArchRise1.angle [°]** | 81.0 | ± | 3.1 | 78.7 | ± | 6.6 | 0.410 | 0.096 |
| **HemalArchRise2.angle [°]** | 78.3 | ± | 2.7 | 77.8 | ± | 4.1 | 0.788 | 0.378 |
| **Diff.Hemal.R1-R2 [°]** | 5.6 | ± | 2.4 | 7.6 | ± | 5.7 | 0.393 | 0.051 |
| **Vertebra.length [µm]** | 0.656 | ± | 0.037 | 0.681 | ± | 0.061 | 0.360 | 0.272 |

### Supplemental Figures

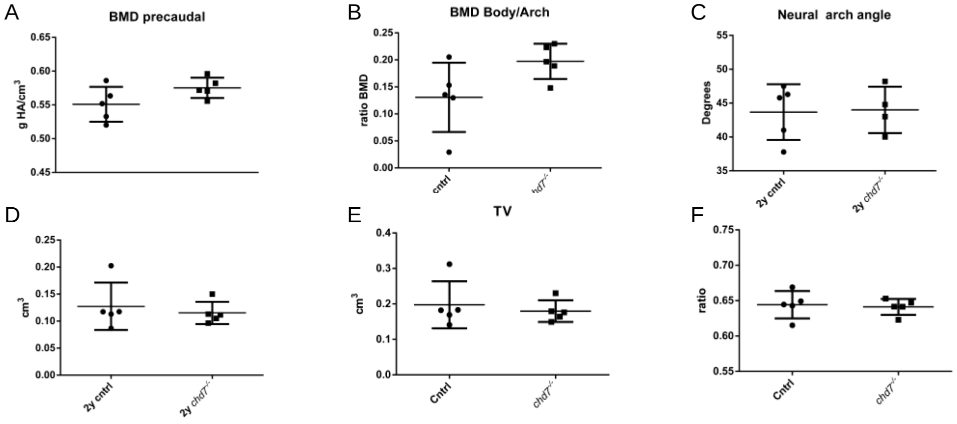


Fig 2. Analysis of key mineralization factors of precaudal vertebrae in 2-year-old *chd7* mutants. A. BMD of whole vertebrae B. ratio of BMD arch/vertebrae body. C. neural arch angle D. BV E. TV and F. ratio BV/TV. Significance of student t-test with Welch's correction is included in the graphs with: *p<0.05; **p<0.01

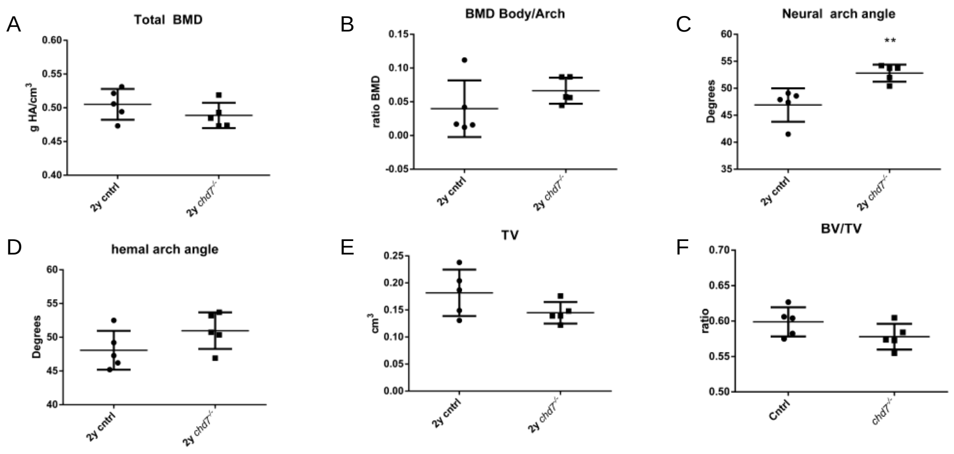


Fig 3 Analysis of key mineralization factors of caudal vertebrae in 2-year-old *chd7* mutants A. BMD of whole vertebrae B. ratio of BMD arch/vertebrae body. C. significant increase in neural arch angle D. hemalvarch angle E. TV and F. ratio BV/TV. Significance of student t-test with Welch's correction is included in the graphs with: *p<0.05; **p<0.01

#### Supplemental Figures

Supplemental Table 4. Statistical analysis of mineralization of the precaudal and caudal vertebrae in 2-year-old zebrafish

| | cntrl 2-year-old | | | CHD7$^{-/-}$ 2-year-old | | | | |
|---|---|---|---|---|---|---|---|---|
| *precaudal vertebra* | n = 5 | | | n = 5 | | | *p(t-test)* | *p(F-test)* |
| **Arch.vBMD [gHA/cm$^3$]** | 0.370 | ± | 0.035 | 0.326 | ± | 0.014 | 0.045 | 0.101 |
| **Body.vBMD [gHA/cm$^3$]** | 0.501 | ± | 0.036 | 0.523 | ± | 0.026 | 0.298 | 0.544 |
| **ArchOpen.angle [°]** | 25.4 | ± | 1.6 | 23.2 | ± | 2.0 | 0.115 | 0.642 |
| **ArchRise1.angle [°]** | 78.4 | ± | 2.4 | 79.7 | ± | 3.6 | 0.558 | 0.440 |
| **ArchRise2.angle [°]** | 76.4 | ± | 1.9 | 77.4 | ± | 4.7 | 0.698 | 0.115 |
| **Diff.R1-R2 [°]** | 3.3 | ± | 3.1 | 4.9 | ± | 6.4 | 0.667 | 0.197 |
| **Body.angle [°]** | 123.6 | ± | 5.7 | 139.8 | ± | 4.1 | 0.002 | 0.603 |
| **Vertebra.length [µm]** | 0.704 | ± | 0.071 | 0.677 | ± | 0.044 | 0.503 | 0.460 |
| *caudal vertebra* | n = 5 | | | n = 5 | | | *p(t-test)* | *p(F-test)* |
| **Arch.vBMD [gHA/cm$^3$]** | 0.498 | ± | 0.033 | 0.464 | ± | 0.020 | 0.096 | 0.333 |
| **Body.vBMD [gHA/cm$^3$]** | 0.538 | ± | 0.028 | 0.531 | ± | 0.019 | 0.657 | 0.443 |
| **NeuralBody.angle [°]** | 124.9 | ± | 9.5 | 124.8 | ± | 2.8 | 0.994 | 0.038 |
| **NeuralArchOpen.angle [°]** | 23.6 | ± | 2.6 | 22.8 | ± | 4.5 | 0.741 | 0.308 |
| **NeuralArchRise1.angle [°]** | 77.8 | ± | 2.8 | 76.0 | ± | 2.5 | 0.317 | 0.803 |
| **NeuralArchRise2.angle [°]** | 79.0 | ± | 2.4 | 81.0 | ± | 3.0 | 0.268 | 0.656 |
| **Diff.Neural.R1-R2 [°]** | 3.1 | ± | 2.4 | 5.0 | ± | 3.4 | 0.327 | 0.517 |
| **HemalBody.angle [°]** | 120.7 | ± | 6.5 | 128.7 | ± | 2.0 | 0.049 | 0.039 |
| **HemanArchOpen.angle [°]** | 24.9 | ± | 5.1 | 24.2 | ± | 2.1 | 0.781 | 0.113 |
| **HemalArchRise1.angle [°]** | 77.6 | ± | 3.9 | 76.4 | ± | 4.4 | 0.657 | 0.825 |
| **HemalArchRise2.angle [°]** | 77.5 | ± | 3.8 | 79.0 | ± | 5.0 | 0.594 | 0.624 |
| **Diff.Hemal.R1-R2 [°]** | 4.6 | ± | 2.3 | 5.5 | ± | 7.1 | 0.791 | 0.049 |
| **Vertebra.length [µm]** | 0.721 | ± | 0.065 | 0.657 | ± | 0.039 | 0.105 | 0.335 |

### Supplemental Figures

Supplemental Table 5. Primer List qPCR

| Target | FW Primer | RV Primer |
|---|---|---|
| *elf1-α* | GTGGCTGGAGACAGCAAGA | AGAGATCTGACCAGGGTGGTT |
| *runx2a* | TGACGTACCTGAGAGGCGT | GCAGCCGTATCCTGCATACC |
| *runx2b* | CGGCTCCTACCAGTTCTCCA | CCATCTCCCTCCACTCCTCC |
| *mgp* | ACGACACGGAGAGAAGTCCT | TGGAGGTGTTTGTGAACCCA |
| *sost* | ACGGACTTATGGAGCCTCAG | TGGAAGGCACTGACCAGAA |
| *ctsk* | TGGAACGGATCAGCAGTGTG | TCTATGCCAACTGACACGGG |
| *acp5a* | GACAGACACGCTGAGCATGAG | ACTGGTTACTTCCTTGAGCTTCCA |
| *postna* | CTGACTCAGCAAAGCAGGTG | GTTCAGTGGCGCAAGTACAG |
| *sp7* | TACCACCGGGAGGTCTTCTT | TTTACCGTACACCTTCCCGC |
| *col2a1a* | CAACACGATGTAGAGGTGGACG | CAGGGTGGCAGAGTTTCAGG |
